# MafK mediates chromatin remodeling to silence IRF8 expression in non-immune cells in a lineage-specific manner

**DOI:** 10.1101/396291

**Authors:** Nitsan Fourier, Maya Zolty, Aviva Azriel, Donato Tedesco, Ben-Zion Levi

## Abstract

The regulation of gene expression is a result of a complex interplay between chromatin remodeling, transcription factors (TFs) and signaling molecules. Cell differentiation is accompanied by chromatin remodeling of specific loci to permanently silence genes that are not essential for the differentiated cell activity. The molecular cues that recruit the chromatin remodeling machinery are not well characterized. IRF8 is an immune-cell specific TF, and thus, serves as a model gene to elucidate the molecular mechanisms governing its silencing in non-immune cells. A high-throughput shRNA library screen in IRF8 expression-restrictive cells enabled the identification of MafK as modulator of IRF8 silencing, affecting chromatin architecture. ChIP-seq analysis revealed three MafK binding-regions (−25kb, −20kb and IRF8 6^th^ intron) in the IRF8 locus. These MafK binding-sites are sufficient to repress a reporter-gene when cloned in genome-integrated lentiviral reporter constructs in expression-restrictive cells only. Conversely, plasmid-based constructs do not demonstrate such repressive effect. These results highlight the role of these MafK binding-sites in mediating repressed chromatin assembly. Furthermore, removal of MafK-int6 binding-region from BAC-IRF8 reporter construct was sufficient to promote accessible chromatin conformation. Taken together, we identified and characterized several MafK binding elements within the IRF8 locus that mediate repressive chromatin conformation resulting in the silencing of IRF8 expression only in non-immune cells.

## Introduction

The regulation of gene expression is not a linear process, but a result of a complex interplay between chromatin modifications, transcription factors (TFs) and signaling molecules. Collectively, this interplay contributes to cell differentiation, maintenance of cell identity, and appropriate responses to the changing environment. Chromatin architecture governs DNA accessibility and, therefore, has a crucial function in establishing, maintaining, and propagating distinct gene expression patterns. Chromatin organization is regulated in part by a complex array of histones post-translational modifications (PTMs), known as the ‘histone code’ (1). Genome-wide studies of such modifications highlighted the correlation of some specific modifications with gene activity. For example, tri-methylation of Histone H3 lysine 27 (H3K27me3) and H3K9me and are associated with repressed chromatin state. In contrast, H3K4me1-3 and histone acetylation (H3ac and H4ac) are associated with transcriptionally active chromatin regions (2). Among the factors that affect the epigenome dynamics are TFs that interact with chromatin and chromatin-modifying enzymes, such as the Polycomb group (PcG) complexes. TFs must gain access to their binding-sites within a chromatin context. However, once bound to DNA, TFs can also modify the chromatin landscape (3).

During the course of cell lineage-specification, cellular gene expression programs need to be adapted to specify the differentiated-cell new identity. Differentiation is accompanied by dynamic changes in chromatin states (4). One of the most studied processes is the differentiation of hematopoietic cells, in which all blood cell-types arise from hematopoietic stem cells (HSCs) through balanced self-renewal and differentiation, and give rise to all mature blood cells. During differentiation, the transcription of genes that are normally expressed in other lineages are often actively suppressed; a crucial but mechanistically not-well characterized hallmark of lineage-instructive TFs (5). One such example is the myeloid TF Interferon Regulatory Factor 8 (IRF8), which plays a key role in myelopoiesis and maintaining the critical balance of the major myeloid subsets (6). Additionally, IRF8 is part of a regulatory complex involved in specification and commitment to B-cell development (7) and controls T-cells functions and activities (8, 9). Aberrant expression of this Interferon-γ (IFNγ) - induced gene (10) is associated with skewed differentiation, which is accompanied by myeloleukemias (11-13). While much is known about the essential mechanisms for its hematopoietic-specific expression, IRF8 silencing in non-immune cells is still uncharacterized. Recently we demonstrated that IRF8 3^rd^ intron controls IRF8 silencing only in non-immune cells (14). This defined element is necessary and sufficient to silence homologous and heterologous gene expression in restrictive cells and possibly acts as a nucleation-core for chromatin remodeling in these cells. Here, we extended our observation using a high-throughput shRNA library screen (15) in IRF8 expression-restrictive cells that enabled the identification of MafK as an IRF8 repressor. MafK belongs to the small Maf (musculoaponeurotic fibrosarcoma) protein family (sMafs), which has vital roles in stress signaling, hematopoiesis, central nervous system function and oncogenesis (16). sMafs can form either homodimers and act as transcriptional repressors (17) or form heterodimers with TFs such as basic leucine zipper (bZIP) proteins and act as repressors or activators, depending on their interacting partner (18, 19). These hetero-complexes bind to Maf response elements (MAREs) (18, 20). Additionally, sMafs form complexes with chromatin modifiers, such as the PcG complexes (21), leading to chromatin remolding and transcriptional activation or repression. Here, we have identified three MafK binding-regions within the IRF8 locus, −25kb, −20kb and IRF8 6^th^ intron, which are sufficient to exert cell-type specific chromatin remodeling even in a random genomic context. Moreover, MafK binding-region in IRF8 6^th^ intron acts as repressive element, affecting the histone PTM signature in the region, and its deletion leads to an open chromatin configuration.

## Results

### High-throughput screening for identifying IRF8 regulator in non-hematopoietic cells

In search of the molecular factors mediating cell-type specific IRF8 silencing, we employed the DECIPHER pooled barcoded lentiviral shRNA libraries, with 5-6 shRNA constructs targeting each of the 9,200 mouse genes (illustrated in Fig. 1a) (15). In brief, this library was transduced to IRF8 expression-restrictive cell-line, and cells harboring shRNAs that led to an increase in IRF8 expression above base-line level, were sorted by FACS and the shRNA construct targeting each cell was identified by its barcode using next generation sequencing (NGS). Specifically, This lentiviral library was transduced to the IRF8 reporter cell-line, NIH3T3, harboring BAC-IRF8.1-GFP construct (NIH3T3-IRF8.1) (14). This genome-integrated BAC harbors 219,907bp encompassing the entire murine IRF8 locus. The BAC-IRF8.1 reporter construct was generated by inserting a reporter cassette containing a fluorescent reporter-gene and an independently transcribed selectable marker at the IRF8 translation start site. This reporter construct authentically reports on IRF8 lineage restrictive-expression in response to IFNγ stimulation; fluorescent in IRF8 expression-permissive macrophage cells and dark in expression-restrictive cells, such as NIH3T3 (murine fibroblast) (14). Following selection for lentiviral transduced cells, the most fluorescent cells were enriched by FACS (top 5%) and as a control, non-fluorescent cells (lower 50%) were collected. Genomic DNA was subjected to high-throughput sequencing to identify the barcode of the integrated shRNAs that were statistically enriched in comparison to the control (Fig. 1b). The final candidates list (Fig. 1c) revealed an over-representation of nuclear proteins (about 50% of the hits), of which close to 30% were TFs (MafK, Fhl2 and Ldb1), further validating our screening approach. Interestingly, the ENCODE database (22) reveals MafK binding in the human IRF8 4^th^ intron (S1 Fig). Additionally, the murine IRF8 3^rd^ intron regulatory element has an *in-silico* predicted MafK DNA binding motif (MatInspector (23)). Together, this placed MafK as our prime candidate for further study.

**Fig 1.**
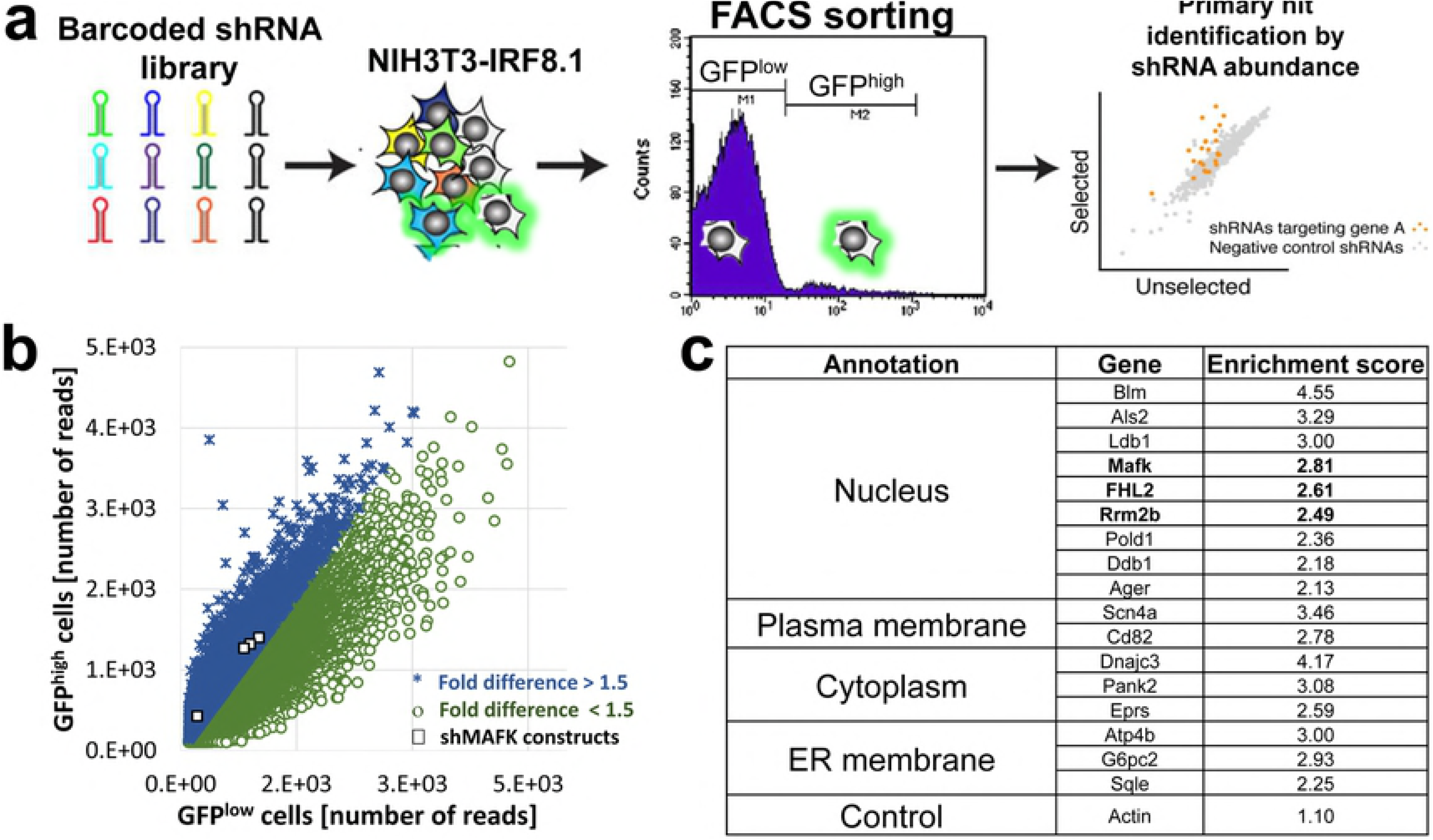
shRNA library screen identifies IRF8 repressor candidates in expression-restrictive cells. (a) Schematic illustration of the DECIPHER shRNA library screen. (b) Scatterplot of number of sequencing reads of each shRNA construct in the GFP^high^ population (y axis) relative to the number of reads in the control GFP^low^ population (x axis) in a representative experiment. For each shRNA construct an enrichment score (the fold-ratio between the sequencing reads in the GFP^high^ population relative to control population) was calculated. shRNA constructs exhibiting enrichment score>1.5 are marked in blue and constructs with enrichment score <1.5 are marked in green. shRNA constructs targeting MafK are marked in black squares. (c) The screen was performed in three replicates and an enrichment score was calculated for each gene, according to the mean score of its shRNA constructs. The table presents the mean±AvDev fold enrichment of the final candidates, which had more than 1.9-fold enrichment score in at least two shRNA constructs for each gene, in all three replicates. The gene candidates were grouped according to their cellular localization annotation. Bold text represents transcription factors. Actin is presented as negative control.

### MafK mediates IRF8 repression in non-hematopoietic cell-line

To validate MafK as IRF8 repressor, MafK was knocked-down (KD) in NIH3T3-IRF8.1 reporter cell-line and the effect on the IRF8-GFP reporter-gene and endogenous IRF8 expression was evaluated by FACS analysis. MafK KD was very effective (down 70%, Fig. 2a) and led to a significant GFP reporter alleviation in IRF8 expression-restrictive NIH3T3-IRF8.1 cells (2.5-fold, Fig. 2b, black bars). Moreover, immunostaining of endogenous IRF8 revealed alleviation of IRF8 expression by more than 2-fold following MafK KD, as evident by FACS analysis (Fig. 2b, grey bars). Furthermore, this FACS analysis clearly demonstrated that in the majority of this cell population, when the reporter-gene expression was alleviated a concomitant alleviation in IRF8 expression was observed (Fig. 2c, right panel), pointing to a direct correlation between GFP reporter-gene expression and endogenous IRF8 expression. Thus, MafK is directly linked to IRF8 repression in restrictive non-hematopoietic cells.

**Fig 2.**
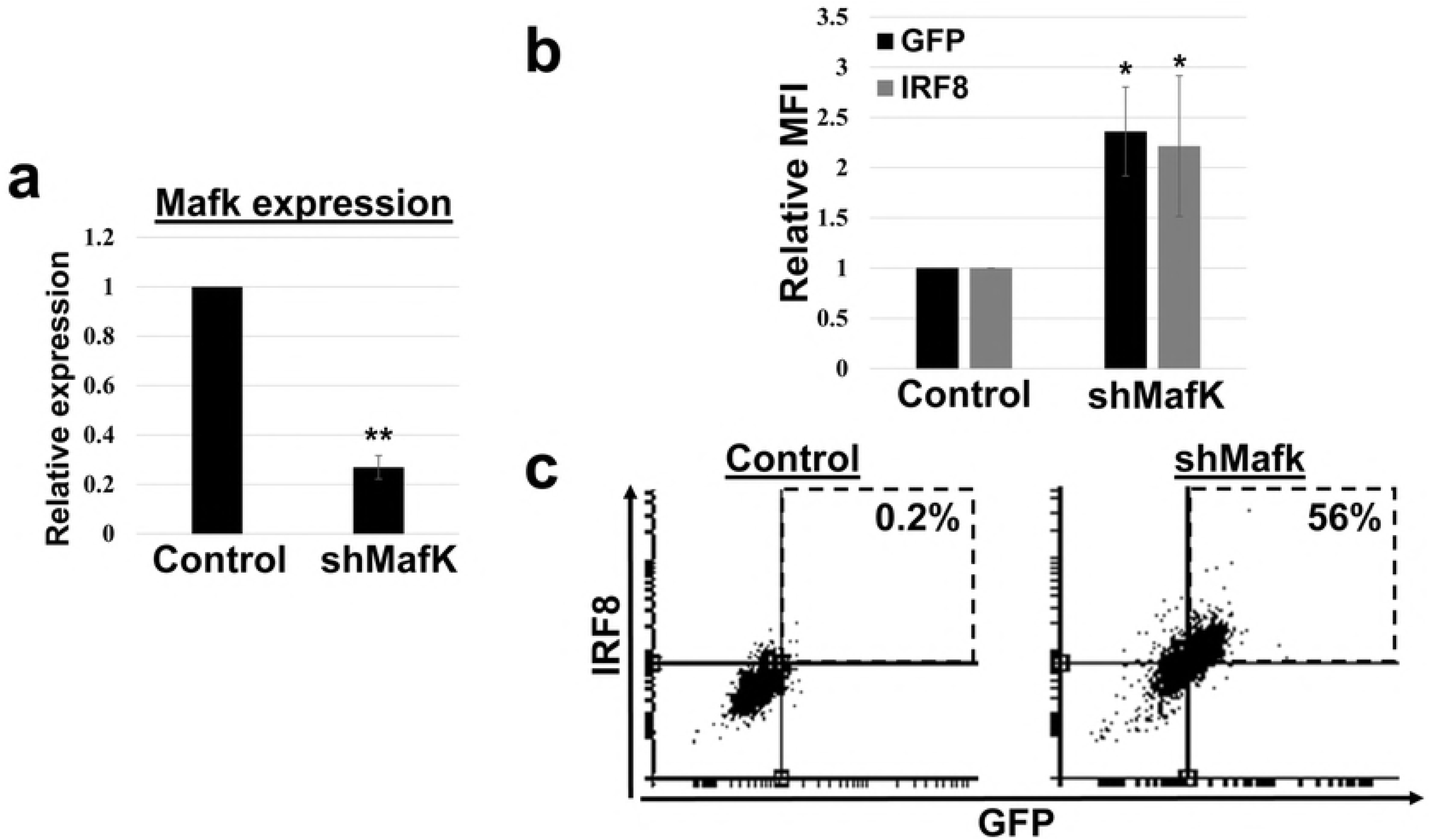
MafK KD is sufficient to alleviate GFP reporter-gene and endogenous IRF8 expression in restrictive NIH3T3 cells. MafK KD was achieved by infecting NIH3T3-IRF8.1 cells with lentiviral vector harboring shMafK or pLKO.1 empty-vector as control. (a) MafK expression level was measured using real-time qRT-PCR. Graph represents mean±AvDev. (b) The cells were immunostained using fluorescence-labeled anti-IRF8 antibodies, and subsequently GFP and endogenous IRF8 expression levels were determined by FACS analysis. Graph represents mean±AvDev of relative GFP (black) and IRF8 (grey) mean fluorescence intensity (MFI) in MafK KD cells. Control cells were determined as 1. (c) Representative FACS dot-plots presenting GFP (x-axis) and IRF8 (y-axis) expression of control (left) and MafK KD (right) NIH3T3-IRF8.1 cells. Red squares represent GFP^high^/IRF8^high^ expressing-cells and their percentage of the total population. *p-value<0.05, **p-value<0.01, student’s t-test, n=3.

To identify the binding sites of MafK within the IRF8 locus, ChIP-Seq analyses were performed in IRF8 expression-restrictive cells (NIH3T3) and IRF8 expression permissive cells (RAW 264.7 (RAW)– murine macrophage cell-line). MafK ChIP-Seq data revealed three distinct binding-sites in IRF8 locus in NIH3T3 and RAW cells (Fig. 3a, black and grey tracks, respectively). IRF8-restrictive (NIH3T3) and permissive cells (RAW) exhibit similar MafK binding patterns; with peaks at −25 kb and −20 kb upstream to IRF8 transcription start site (TSS) and within the IRF8 6^th^ intron (MafK25, MafK20 and MafK-int6, respectively). While differential binding pattern between restrictive and permissive cells was expected, the results reflects the reported MafK dual activity, as an activator or repressor, depending on its interacting partner (16, 18, 20). A JASPAR database search (24) predicts MafK DNA binding motifs with varying affinity scores in each of the ChIP-Seq binding-regions. The ChIP-Seq data was further validated by ChIP-PCR, enriching MafK bound DNA from the above-mentioned three binding-sites (S2 Fig). Although MafK binding profile revealed similar MafK occupancy in NIH3T3 and RAW cells, differential enrichment analysis (Fig. 3b) revealed highly significant (p-value of 7.10^−11^) 4-fold MafK binding enrichment in the −25 kb region in RAW cells in comparison to NIH3T3 cells. The −20 kb region also exhibited significant 2-fold enrichment (p-value of 9.10^−5^) in RAW cells, while IRF8-int6 showed no differential enrichment. The significant difference in MafK binding, favoring IRF8 expression-permissive cells, suggests that MafK25 and MafK20 sites act as activators. In order to extend these results, the regulatory functionality of each binding-region was further investigated using various reporter assay systems, as discussed hereafter.

**Fig 3.**
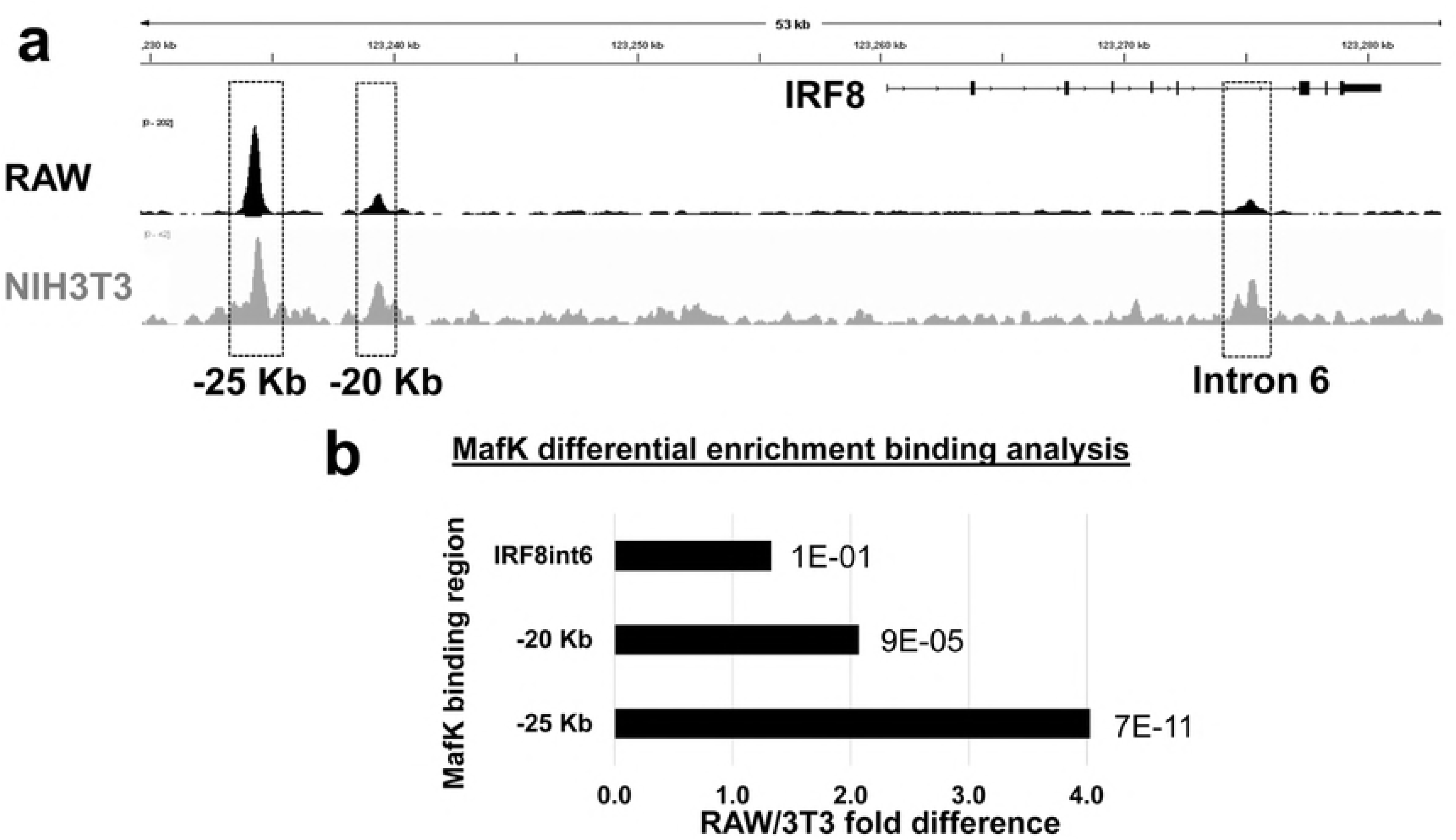
MafK binding profile in IRF8 locus in IRF8 expression-restrictive and permissive cells. (a) Representative tracks from RAW (top, black track) and NIH3T3 (bottom, grey track) MafK ChIP-seq. Black rectangular represent three MafK binding-regions: −25 kb and −20 kb upstream to IRF8 TSS and IRF8 6th intron. (b) Differential enrichment analysis of MafK binding-regions in RAW vs NIH3T3 cells. Graph presents fold difference of RAW/NIH3T3 calculated from three replicates. p-value of each ratio is also given.

MafK was reported as an interacting factor with chromatin-modifying complexes leading to repressed chromatin conformation (21, 25). Thus, the involvement of MafK in mediating repressed chromatin, characterized with enriched H3K27me3 deposition, was examined at the IRF8 locus. To this end, H3K27me3 ChIP-qPCR was performed on NIH3T3-IRF8.1 cells that were KD for MafK. As evident in Fig. 2, effective KD was accompanied by elevated GFP reporter-gene and endogenous IRF8 expression. Thus, cells were FACS-sorted according to their GFP expression and ChIP-qPCR for H3K27me3 modification was performed on the GFP^high^ population (effective MafK KD) and GFP^low^ population (non-effective MafK KD), as control. H3K27me3 ChIP-qPCR demonstrated that MafK KD was accompanied by a significant decrease in H3K27me3 repressive modification over the reporter-gene (Fig. 4a) as well as over the endogenous IRF8 (Fig. 4b), resulting in a less-condensed chromatin conformation accompanied by alleviation of gene expression (Fig. 2b). These results underscore the effect of MafK on chromatin remodeling, as evident by H3K27me3 deposition, as part of its regulatory mechanism in silencing IRF8.

**Fig 4.**
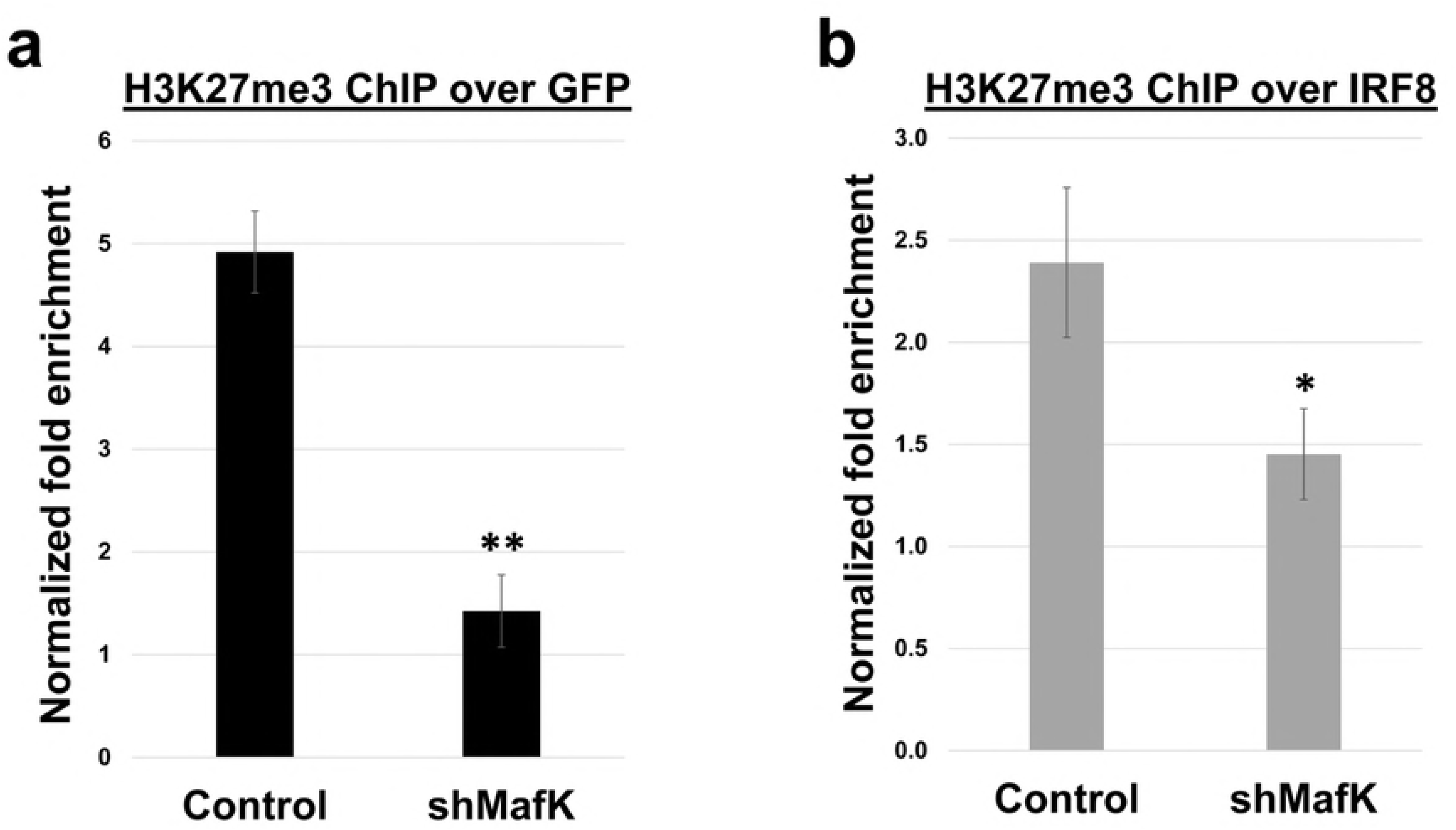
MafK affects deposition of repressive H3K27me3 PTM over IRF8 locus. NIH3T3-IRF8.1 cells were infected with shMafK lentiviral vector or pLKO.1 empty-vector as control. Infected cells were sorted using FACS, according to GFP expression, into MafK KD (GFP^high^) and control cells (GFP^low^) and subjected to H3K27me3 modification ChIP-qPCR using primers targeting either the GFP reporter-gene (a) or the endogenous IRF8 (b). Graph represents the normalized fold enrichment ratio of Ab enrichment vs non-specific IgG Ab. * p-value<0.05, **p-value<0.01 student’s t-test, n=2.

To further delineate the role of Mafk in silencing IRF8 in expression-restrictive cells in comparison to expression-permissive cells, we investigated the effect of each binding-region (MafK25, MafK20 and MafK-int6) on the IRF8 promoter in a reporter gene assays. On one hand, MafK may act as a transcriptional repressor or activator (16, 18, 20), depending upon its interacting partner. On the other hand, MafK recruits chromatin modifiers that may render the IRF8 locus accessible or inaccessible to the transcriptional machinery (21, 25) in a cell-type specific manner. Luciferase reporter assays were performed employing two strategies that may shed light on the mode of MafK action on IRF8 expression in restrictive or permissive cells. In the first strategy, we used transiently expressed Luciferase reporter cassettes cloned in a plasmid (pGL3, see illustration in Fig. 5a), which does not integrate into the genome and thus does not assemble chromatin conformation (26). Therefore, this type of assay allows to determine the direct effect of TFs interacting with MafK binding-sites, regardless of chromatin state. In the second strategy, we used the same reporter cassettes cloned in a retroviral vector, (pMSCV, Fig. 5a). This retroviral vector randomly integrates into infected cell genome and assembles chromatin conformation. Consequently, the chromatin environment effects on the reporter-gene expression. In this reporter system the effect of each MafK binding-site on chromatin assembly is measured. Thus, these two reporter systems may point to different regulatory mechanisms. Specifically, each of MafK binding-regions was PCR-amplified and cloned upstream to IRF8 promoter driving the expression of the Luciferase reporter-gene and subsequently cloned to the plasmid pGL3 or to the retroviral vector pMSCV (as illustrated in Fig. 5a). These reporter constructs, as well as control empty-vector, were transfected/infected to NIH3T3 cells (IRF8 expression-restrictive cells) and to RAW cells (IRF8 expression-permissive) and Luciferase levels were measured. No change in reporter-gene activity, in either of MafK reporter constructs, was observed when using the transient Luciferase reporter plasmid system in expression-restrictive NIH3T3 cells (Fig. 5b, left panel). Similarly, only minor changes in Luciferase levels were noted when using the transient reporter system in IRF8-permissive RAW cells; while MafK25 region exhibited slight reduction in reporter activity, MafK-int6 exhibited slight increase in reporter activity (Fig. 5b, right panel). Conversely, Luciferase activity was significantly reduced with all reporter constructs when using the retroviral Luciferase system only in expression-restrictive cells (∼65%, compare Fig. 5c, left and right panels). It is clear that in expression-permissive cells reporter-gene activity was not reduced when using the retroviral reporter system. On the contrary, MafK25 and MafK20 regions exhibited increased reporter activity, albeit not statistically significant (Fig. 5c, right panel). Collectively, MafK binding-sites exhibit cell-type specific repression that probably reflects cell-type specific chromatin remodeling.

**Fig 5.**
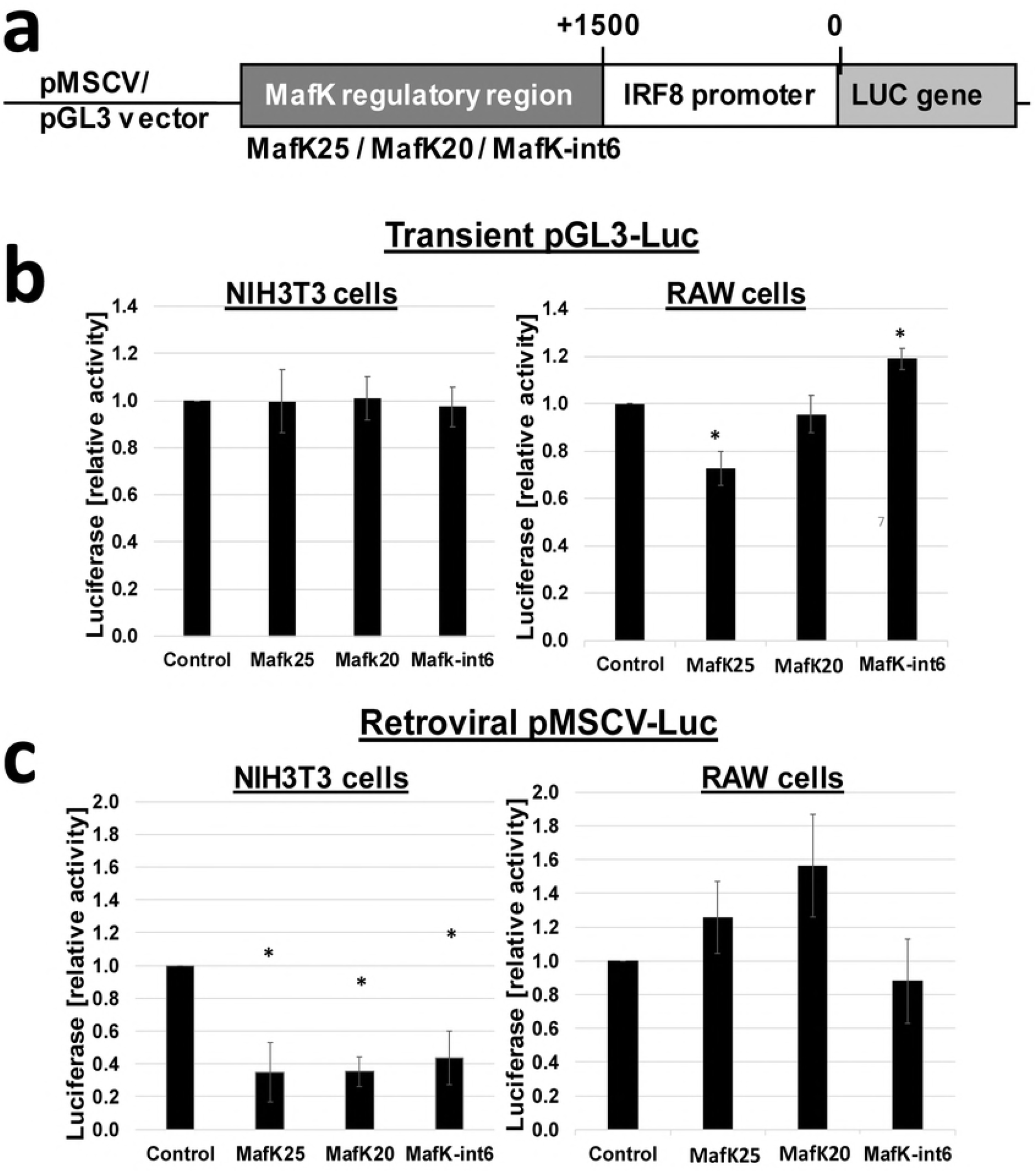
MafK mediates chromatin remodeling in IRF8 locus in a cell-type specific manner. (a) Illustration of MafK-Luciferase reporter constructs. pGL3 (plasmid) or pMSCV (retroviral) vectors were used as backbone to create MafK-Luciferase reporter constructs. MafK binding-regions located −25 kb (MafK25), −20 kb (MafK20) and IRF8int6 (MafK-int6) were each separately cloned upstream to IRF8 promoter. (b) Expression-restrictive NIH3T3 (left panel) and permissive RAW (right panel) were transfected with transient pGL3-IRF8p-Luc-MafK constructs. Transient Luciferase activity was normalized to Rennila luciferase activity and total protein amount. (c) Expression-restrictive NIH3T3 (left panel) and permissive RAW (right panel) were infected with retroviral vectors pMSCV-IRF8p-Luc-MafK. Retroviral Luciferase activity was normalized to vector genomic copy number, as determined using real-time qPCR, and total protein amount. Relative Luciferase activity was calculated as ratio between MafK construct activity and control (Luciferase empty-vector). *p-value<0.05, student’s t-test, n=3-4.

Next, Luciferase-MafK reporter experiments combined with MafK overexpression were conducted to directly link MafK to the silencing effect exerted by MafK binding-regions in NIH3T3. For that purpose NIH3T3 and RAW cells were transfected with the above-mentioned transient Luciferase-MafK reporter plasmids, together with pMSCV-MafK overexpression vector, and MafK overexpression was verified by real-time qPCR (**Error! Reference source not found.**). MafK overexpression in expression-restrictive NIH3T3 cells resulted in approximately 35% reduced reporter activity in all reporter constructs (Fig. 6a). In RAW cells strong reduction (66%) was noted only in MafK25 construct (Fig. 6b). Collectively, these results point to a direct involvement of MafK in the repressive regulatory activity exhibited in expression-restrictive cells by MafK binding-regions.

**Figure 6.**
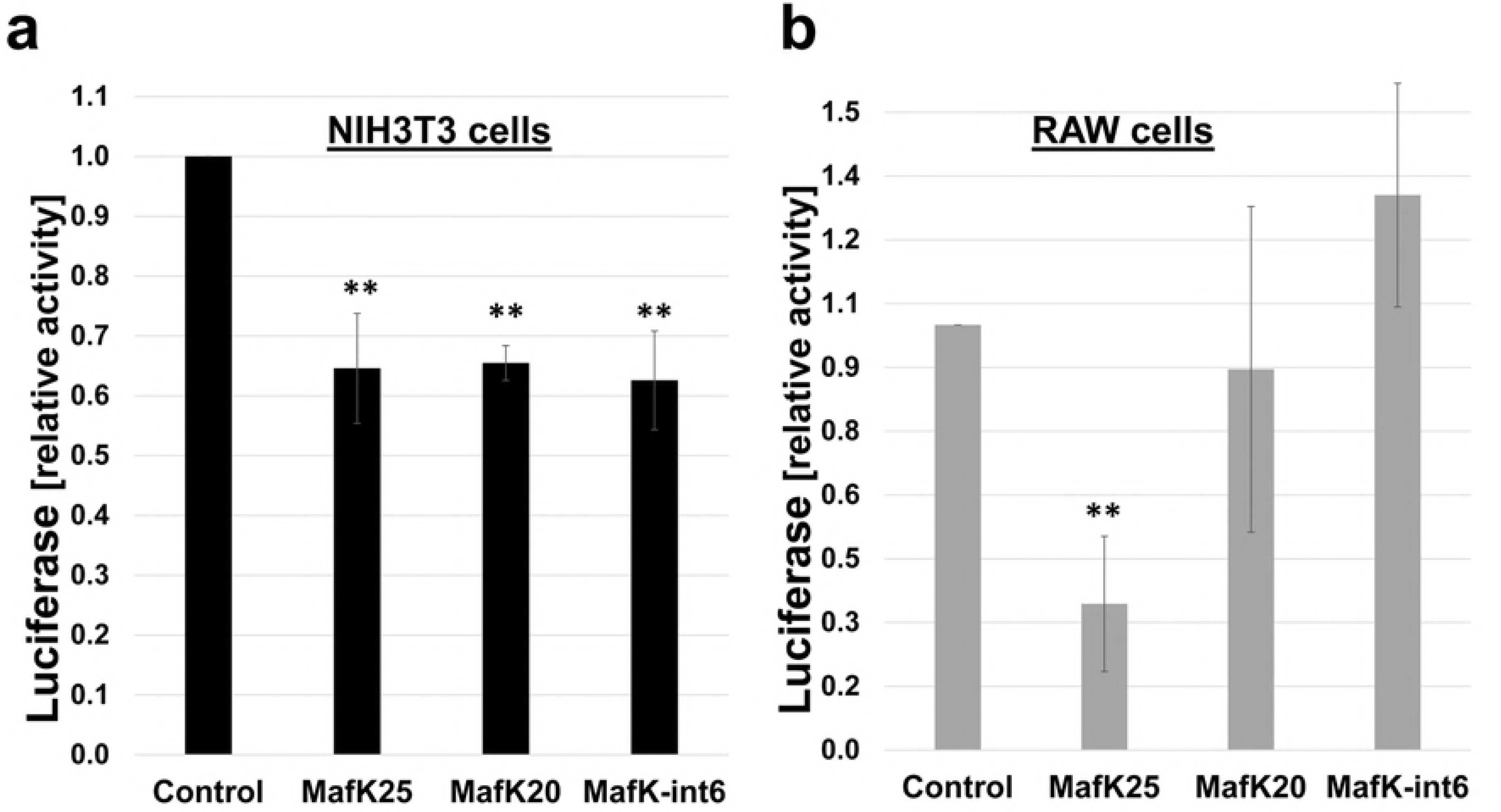
MafK mediates the repressive activity of MafK regulatory-regions in expression-restrictive cells. NIH3T3 (a) or RAW (b) cells were transiently transfected with pGL3-IRF8p-Luc-MafK reporter constructs and pMSCV-MafK overexpression vector. Luciferase activity was normalized to Rennila activity and total protein amount. Relative Luciferase activity was calculated as ratio between MafK construct activity and control (Luciferase empty-vector). **p-value<0.01 student’s t-test, n=3-4.

Each of MafK binding-regions harbors several Maf response elements (MAREs) with different affinity scores (JASPAR database (24)), among them a MafK/Bach1 composite element, a known co-repressor of MafK (27, 28). As our results in Fig. 5 point to the role of MafK as mediator of chromatin conformation, the functionality of each the MAREs was characterized using the retroviral Luciferase reporter assay in expression-restrictive NIH3T3 cells. Each region was divided into segments according to the distribution of putative binding-sites, and cloned separately upstream to the IRF8 promoter in the pMSCV-IRF8p-Luc reporter construct. NIH3T3 cells were infected with individual segmented constructs and relative Luciferase reporter activity was calculated. MafK25- This binding-region has numerous putative MAREs, of which three exhibit high affinity score (>10), including a MafK/Bach1 composite element. The MafK25 region was divided into three segments (A-C, Fig. 7ai): (A) harboring a MafK/Bach1 element and two lower-score MAREs (B) harboring two high-score MAREs and one low-score MARE (C) having no MARE (control). As expected, the Mafk25 segmentation-analysis (Fig. 7aii) demonstrated decreased Luciferase activity only in constructs harboring MAREs (MafK25_A and MafK25_B). MafK25_C, having no MafK DNA binding motif, had no effect on reporter activity, similar to empty-vector. MafK20 has only two putative MAREs; a low score motif (5) and a MafK/Bach1 composite element, having the highest score (17) of all MAREs in all of MafK regions.

**Fig 7.**
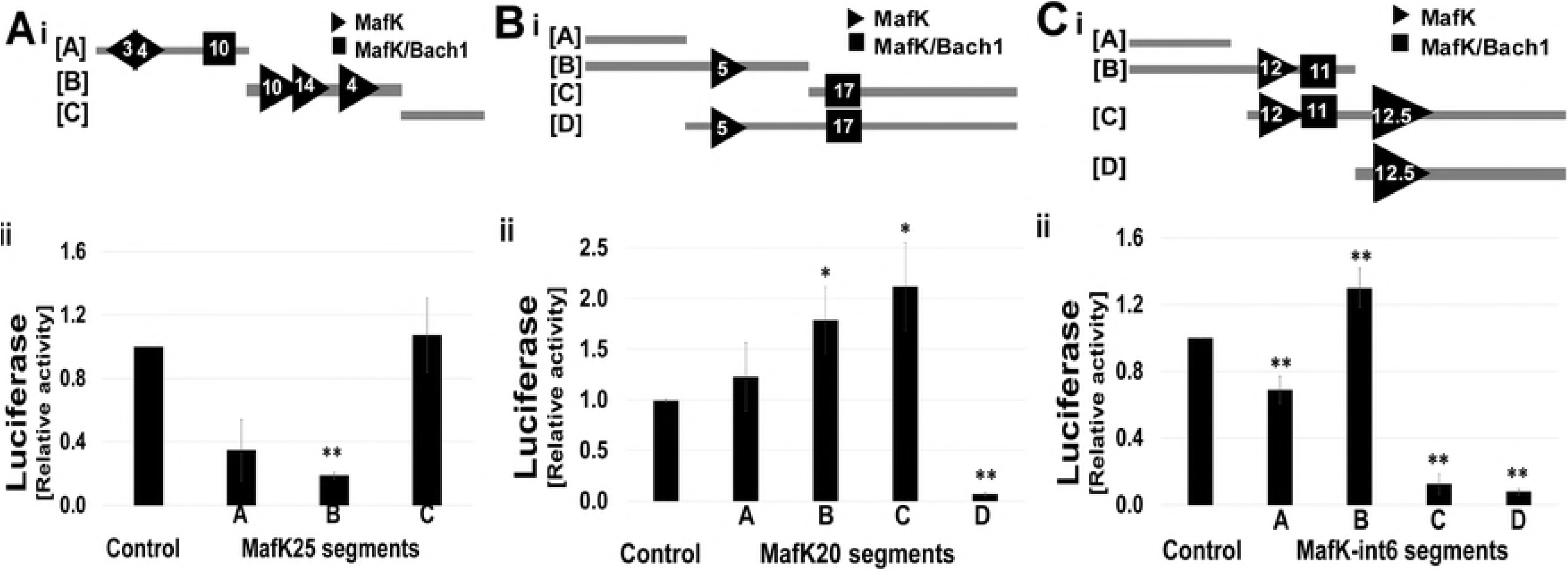
Differential MARE activity in MafK regulatory-regions. MafK25, MafK20 and MafK-int6 regulatory-regions (ai,bi,ci respectively) were divided into segments and MARE sites are illustrated in triangular (MafK) and rectangular (MafK/Bach1) with their predictive affinity score, as determined by the Jasper motif discovery (24). NIH3T3 were infected with pMSCV-IRF8p-Luc construct harboring either MafK25, MafK20 or MafK-int6 segmented regulatory-regions (aii,bii,cii respectively). Retroviral Luciferase activity was normalized to vector genomic copy number, as determined using real-time qPCR. Relative Luciferase activity was calculated as ratio between MafK construct activity and empty-vector (control). *p-value<0.05, **p-value<0.01 student’s t-test, n=2-3.

MafK20- This binding-region was divided into four segments: (A) having no MARE (control) (B) harboring the low-score MARE (C) harboring the MafK/Bach1 element (D) harboring both MAREs (Fig. 7bi). As evident in Fig. 7bii, control segment (MafK20_A) exhibited no significant change in reporter activity relative to empty-vector. MafK20_B also showed no decrease in Luciferase activity, but rather a slight increase. However, since this segment harbors a low-score MARE, this segment was expected to not convey significant repressive activity. Surprisingly, segment C, harboring a high-score MafK/Bach1 element, also did not exhibit repressive activity, but rather significant 2-fold increase of reporter activity. However, segment D, harboring both MAREs, elicited more than 90% decrease in reporter activity. Taken together, only cooperative effect between the two MafK binding-sites lead to reporter repression.

MafK-int6- This binding-region harbors three high-score MAREs (11-12.5) in close proximity to each other (∼50 bps), one of which is a MafK/Bach1 composite element. The MafK-int6 region was divided into four segments (Fig. 7ci): (A) having no MARE (control) (B) harboring two MAREs, one of which is a MafK/Bach1 element (C) harboring all of MAREs (D) harboring the highest score MARE. In MafK-int6 binding-sites segmentation analysis (Fig. 7cii), control segment (MafK-int6_A) exhibited slight but significant decrease in reporter activity relative to empty-vector. Although MafK-int6_B harbors two high-score MAREs, it did not exhibit decreased Luciferase activity, but rather a slight increase. Segment C, harboring an additional MARE with the highest score in this region (12.5), exhibited a significant decrease in reporter activity. Finally, segment D, harboring only this high score MARE, exhibited similar decrease (90%) in Luciferase activity. The data suggest that the main repression activity exerted by MafK-int6 region is elicited by the MARE element located in segment D, which counteracts the activity of the other segments.

Collectively, the deletions analysis within the three MafK binding-regions (Fig. 7) further emphasize the fact that MafK is directly involved in suppressive regulation of MafK binding-regions, as control segments, harboring no MafK motif, show similar reporter activity as control empty-vector.

### MafK-int6 regulatory region mediates repression via chromatin remodeling in IRF8 locus in BAC reporter system

Previously, we demonstrated that IRF8 harbors a regulatory region in its 3^rd^ intron, mediating restrictive-expression via chromatin remodeling in non-hematopoietic cells. Deletion of IRF8 3^rd^ intron in the BAC-IRF8.1 construct resulted in alleviation of GFP reporter-gene expression in restrictive NIH3T3-IRF8.1-▽int3 cells (14). Based on the above results, we propose that MafK-int6 binding-region also acts as an intronic regulatory element modulating chromatin conformation. To this end, the BAC-IRF8.1-GFP reporter system was employed. As mentioned above, the BAC harbors all of IRF8 regulatory regions and regains original chromatin architecture regardless of integration point (29, 30). A new BAC-IRF8.1-▽MafK-int6 was generated by replacing the MafK-int6 binding-region with a Zeocin resistance cassette in bacteria, as previously described (14). Subsequently, IRF8-restrictive NIH3T3-IRF8.1-▽MafK-int6 reporter cell clones were generated by transfecting this BAC construct as described under Methods. MafK-int6 binding-region repressive activity in expression-restrictive NIH3T3 cells was analyzed by FACS. Since IRF8 is IFNγ inducible gene only in expression-permissive cells (14), the level of the reporter gene was also tested in the presence of IFNγ. The data clearly showed that deletion of MafK-int6 binding-region resulted in alleviation of GFP reporter expression relative to the control (complete BAC-IRF8.1) with or without IFNγ induction (Fig. 8a). Specifically, 1.4-fold GFP alleviation was observed even without IFNγ induction, which was further enhanced by IFNγ treatment, resulting in 3-fold induction of GFP expression in comparison to control (Fig. 8b, black and grey bars respectively). As expected, analysis of endogenous IRF8 mRNA expression level in these cells showed no such alleviation since the endogenous 6^th^ intron is intact (Fig. 8c). These results highlight the role of this intron in IRF8 regulation in expression-restrictive cells.

**Fig 8.**
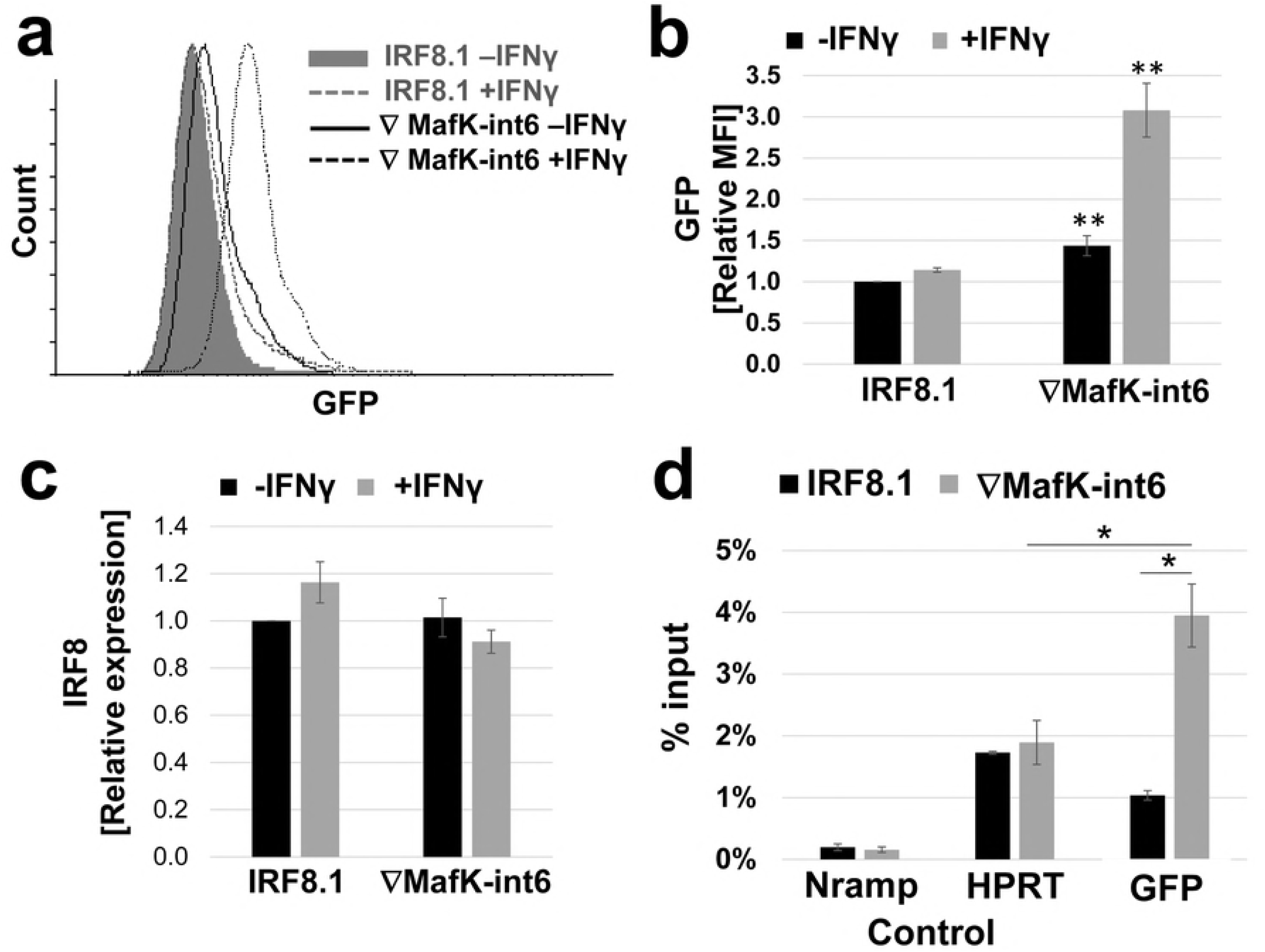
MafK-int6 regulatory-region mediates repressive histone PTM signature in restrictive NIH3T3-IRF8.1 reporter cells. NIH3T3 clones harboring BAC-IRF8.1 (control) or BAC-IRF8.1-▽MafK-int6 were either treated of not-treated with IFNγ for 16 hrs. Cells were than taken for FACS analysis of GFP reporter expression level. (a) Representative histogram of NIH3T3-IRF8.1 (untreated, grey-filled histogram, and IFNγ-treated, dotted-grey histogram) and NIH3T3-▽MafK-int6 (untreated, black histogram, and IFNγ-treated, dotted-black histogram) cells. (b) Graph shows relative mean fluorescence intensity (MFI) of treated and untreated NIH3T3-IRF8.1 and NIH3T3-▽MafK-int6 cells. Untreated NIH3T3-IRF8.1 were determined as 1. Results are mean±AvDev of two NIH3T3-▽MafK-int6 clones, n=3. (c) Endogenous IRF8 mRNA expression level was measured using real-time qRT-PCR. Results are mean±AvDev of two NIH3T3-▽MafK-int6 clones, n=3. (d) NIH3T3 cells harboring either BAC-IRF8.1 or BAC-IRF8.1-▽MafK-int6 were subjected to H3K27ac ChIP-qPCR. Nramp1 promoter serves as negative control and HPRT promoter serves as positive control. Enrichment over the GFP gene was detected using primers targeting GFP coding region. Value are mean±AvDev of %input. *p-value<0.05, student’s t-test, n=2-3.

The data presented in Fig. 4 showed that MafK KD in NIH3T3-IRF8.1 cells resulted in decreased repressive H3K27me3 histone PTM enrichment over the GFP reporter-gene as well as the endogenous IRF8 gene. Accordingly, we expected that deletion of MafK-int6 repressor element will result in a change in the chromatin landscape, *i.e.* open chromatin conformation. Therefore, H3K27ac (a common marker for open chromatin (31)) ChIP-qPCR was performed across the GFP reporter-gene in cells harboring the BAC constructs, BAC-IRF8.1-▽MafK-int6 and control BAC-IRF8.1 (Fig. 8d). Endogenous HPRT promoter (expressed house-keeping gene) and endogenous Nramp1 promoter (repressed hematopoietic gene) regions were used as positive and negative controls, respectively. In NIH3T3-IRF8.1-▽MafK-int6 cells the H3K27ac PTM enrichment was 4-fold higher over the GFP reporter region compared to control NIH3T3-IRF8.1 cells. Moreover, a 20-fold enrichment was observed compared to the negative (repressed) control promoter. Even compared to the positive control, H3K27ac enrichment over GFP reporter-gene was significantly higher (2-fold). As expected, each control gene exhibited similar PTM enrichment for both BAC harboring cell-types. Additionally, the positive control, HPRT exhibited approximately 10-fold H3K27ac enrichment relative to the repressed Nramp1 promoter. Collectively, analysis of active (H3K27ac) histone PTM points to a more accessible chromatin-state over the GFP reporter-gene on the BAC construct following removal of MafK-int6 regulatory-region in IRF8 expression-restrictive NIH3T3 cells. Since we demonstrated that the BAC-IRF8 reporter constructs authentically report on IRF8 endogenous gene expression (Fig. 2d and (14)), it strongly suggests that multiple intronic elements, the 6^th^ and the 3^rd^ introns, orchestrate IRF8 chromatin architecture in expression-restrictive cells.

## Discussion

Lineage-specific gene expression is a two-facet process; while one facet is involved in regulating cell-type specific expression, the other executes an opposing program aimed at long-term gene silencing in expression-restrictive cells. Chromatin architecture is the determining factor in this cell-fate decision process. While it is clear that condensed chromatin is essential for sustained repression, the molecular cues recruiting the chromatin remolding machinery are not well characterized. Specifically, the roles of the TF IRF8 in hematopoiesis are well documented (7, 32-34). Yet, the molecular mechanisms preventing misexpression in restrictive cells are not characterized. Recently, we reported that IRF8 3^rd^ intron is sufficient and necessary to initiate gene silencing in non-hematopoietic cells, highlighting its role as a nucleation core for repressed chromatin during differentiation (14). The current study was aimed at identifying DNA binding factors that may serve as the mediators or cues that recruit the chromatin remodeling machinery. To this end, a high-throughput lentiviral shRNA library was screened in a reporter cell-line, which led to the identification of MafK as an IRF8 repressor in expression-restrictive cells (14). MafK acts as repressor by forming homo- or hetero- dimers with other TFs, mainly with Bach1 (17, 28, 35). Further, MafK has been shown to interact with histone modifying complexes, such as the PcG complexes, through its partner MAT2α, mediating repressive histone modifications (21, 25). MafK was established as a bona-fide IRF8 repressor, as MafK KD in expression-restrictive GFP-IRF8 reporter cells resulted in significant GFP reporter alleviation (Fig. 2b and c). Moreover, MafK directly affected endogenous IRF8 expression, as IRF8 protein expression was also alleviated following MafK KD (Fig. 2b and c). This co-elevated expression of GFP and IRF8 following MafK silencing underscores the authenticity of the BAC-IRF8-GFP reporting system. MafK ChIP-Seq in IRF8-restrictive (NIH3T3) and permissive (RAW) cells revealed three MafK binding-sites in IRF8 locus: −25 kb, −20 kb and MafK-int6, respectively (Fig. 3b and c). These relatively “low-resolution” binding-regions (∼200bp) are abundant with putative MafK response elements (MAREs) (Fig. 7ai, bi and ci). The ubiquitous MafK expression (S4 Fig.) and binding profile between expression-restrictive and permissive cells is not surprising due to MafK dual activity, depending on its interacting partner (18). Yet, a higher degree of MafK occupancy in permissive cells compared with restrictive cells was noted for binding-region −25kb (p-value= 7.10^−11^) and to a lesser extent for −20kb (p-value=9.10^−5^) but not for IRF8-int6 (p-value=10^−1^) (Fig. 3b).

Individual MafK binding-regions exhibited cell-type specific effect using lentiviral reporter system, *i.e*. significant reduction in Luciferase reporter-gene activity in IRF8 expression-restrictive cells as opposed to enhanced activity in IRF8 expression-permissive cells (Fig. 5c). Moreover, these elements were sufficient to exert chromatin remodeling in a random genomic context. This lineage-specific regulatory effect was observed only when the reporter vectors were integrated into the genome and assembled chromatin conformation. This did not take place in transient transfection assays, where the transfected plasmid DNA does not assemble proper chromatin structure (Fig. 3c and d, black columns) (26). Further, MafK-int6 region acts as an intragenic *cis*-repressor as deletion of this region from the BAC-IRF8 construct in expression-restrictive cells was sufficient to alleviate the expression of the GFP reporter-gene. As expected, no effect was observed on the expression of the endogenous IRF8 in the same cells (Fig. 8c). Moreover, the deletion of MafK-int6 region was accompanied by transition towards accessible chromatin conformation, evident by significant increase in H3K27ac histone mark in this region (Fig. 8d). Finally, MafK KD coincided with reduced repressive H3K27me3 histone PTM deposition (Fig. 4). Together, our results underscore the role of MafK in silencing IRF8 expression by affecting local chromatin architecture. Similar silencing elements have been previously described by Jermann et al. (36), demonstrating that small CpG-islands can autonomously recruit PcG components and establish an H3K27me3 domain, suggesting that local sequence autonomy may be a general feature of PcG recruitment in the mouse genome. We propose that in non-hematopoietic cells MafK binding-regions serve as recruiting elements for the repressive PcG complex, leading to heterochromatin formation and subsequent IRF8 silencing. This is supported by previous reports of MafK interactions with repressive chromatin modifying complexes, such as the NuRD and PcG complexes, regulating downstream gene *in*-*cis* (21, 37). Interestingly, while in restrictive cells all regions exhibited repressive reporter activity, in permissive cells only MafK25 and MafK20 enhanced reporter-gene expression. Concomitantly, these regions also exhibited significantly higher degree of MafK occupancy in permissive cells compared with restrictive cells (Fig. 3b). This may be due to the fact that the MARE motif is shared with the AP-1 TF family proteins, such as Jun and Fos, which are known MafK binding-partners (20, 35). Jun and Fos are also known IRF8 activators, part of the IFNγ signaling pathway (38), and exert cell-type regulatory activity (39). Alternatively, other members of the Maf family, c-Maf or MafB, which are highly expressed in IRF8-permissive cells, may preferentially bind the MARE sites and compete with MafK binding (40, 41), leading to transcriptional activation. The dual-functionality of a single *cis-*regulatory element suggests a mechanism for efficient toggling between effector states in changing environments. This dual-regulatory phenomena has been described in developmental enhancers and lineage-specific genes controlling cell-identity plasticity (37, 42, 43).

MafK exhibited preferential binding for specific MARE sites in each of the regulatory binding-regions in IRF8-restrictive cells. Only regulatory segments harboring a high-score MARE site exhibited repressive effect, as was evident by retroviral-mediated reporter assay, as opposed to low-score MAREs and/or MafK/Bach1 composite element (Fig. 7). This is expected as silencing elements specificity is finely tuned, often being affected by flanking sequences and *cis*-elements (44). The fact that elements not harboring a MARE motif did not elicit repressive reporter-gene activity in IRF8-restrictive cells (Fig. 7) highlights the direct involvement of MafK in suppressive regulation of MafK binding-regions.

Interestingly, some MAREs exhibited enhanced reporter activity when separately present in the reporter segment (MafK20_B, MafK20_C and MafK-int6_B, Fig. 7), but when combined with an additional MARE site demonstrated significant silencing effect (MafK20_D, MafK-int6_C and MafK-int6_D, Fig. 7). Cooperative binding of a TF to multiple DNA binding-sites can intensify or modify its repressor function (44). Such cooperative effect was also suggested for the mouse HoxD PcG response element (PRE), while the full-length PRE exhibited substantial silencing effect, two of its fragments enhanced gene expression and one fragment demonstrated a silencing effect (45).

Lastly, as Bach1 is the most studied MafK co-repressor, we examined its possible role in MafK mediated silencing of IRF8. Although all of MafK binding-regions harbor a MafK/Bach1 composite DNA binding motif, segmented element harboring this motif did not exert repressive reporter gene activity (Fig. 7). Bach1 overexpression combined with transient MafK Luciferase reporter assay also did not significantly reduce reporter gene activity (S5 Fig.). Moreover, Bach1 ChIP-qPCR did not exhibit binding in either of MafK regulatory regions (S6 Fig.). Thus MafK`s repressive partner in IRF8-restrictive cells is yet an unidentified repressor, or that MafK silencing activity is a result of MafK homodimerization (17).

Taken together, we propose that MafK binding-regions serve as dual-functioning regulatory elements. In IRF8-restrictive cells, these sites are occupied by MafK homo- or hetero- dimers, which contributes to the repressive regulatory equilibrium. Conversely, in IRF8-premisive cells, these regions may preferentially be occupied by other Maf (46, 47) or AP-1 family members (38), thereby competing with the repressive MafK complex, shifting the equilibrium to an active transcriptional state. Additionally, we suggest that the molecular mechanisms preventing misexpression of lineage-specific genes is composed of several checkpoints. Thus far, we have identified two such control checkpoint elements: the IRF8 3^rd^ intron (14), acting also as a transcriptional activator in expression-permissive cells (Khateb et al, unpublished data), and the multiple MafK regulatory binding-sites that also function in a dual manner.

## Methods

### Cell-lines

NIH3T3 (Mouse embryo fibroblast), RAW (RAW267.4, Murine monocytes/macrophages-like) and 293FT (Human embryonal kidney) were obtained from ATCC, Manassas, Virginia, USA (CRL-1658, TIB-71 and CRL-3216, respectively). These cell-lines were maintained in DMEM supplemented with 10% FCS, 2.5 μg/ml Amphotericin and 50 μg/ml Gentamycin Sulfate (Biological Industries, Beit-Haemek, Israel).

### Pooled shRNA library

The mouse DECIPHER (15) pooled shRNA library screen was performed according to the DECIPHER user manual. To produce library viral pool, 120 µg of library DNA in combination with lentiviral packaging plasmids were transfected to 293FT cells in ten 15 cm tissue culture plates using Lipofectamine2000 and PLUS transfection reagents (Invitrogen, Carlsbad, CA, USA). The viral titer was estimated using the TagRFP reporter-gene on the shRNA library vector. The MOI was determined from the calibration curve provided by the library’s user manual.

For the library screen, 2.10^6^ NIH3T3-IRF8.1 cells were seeded on the day of infection in a 15 cm tissue culture plate, in three replicates. Cells were transduced at an MOI of 0.7 to ensure infection of no more than a single shRNA construct in each cell. 72 hrs post-infection, Puromycin selection was added. 48 hrs post-selection, the cells were collected, counted and ∼3.10^7^ cells (>1000-fold library complexity) were taken for FACS sorting. The top 5% GFP^+^ cells were sorted as positive population and the 50% GFP^−^ cells were sorted as negative (control) population. Following the sorting process, genomic DNA was extracted from each population by Phenol:Chloroform extraction followed by Ethanol precipitation. The DNA was diluted at estimated 2 µg/µl concentration, assuming yield of 10 µg DNA per 10^6^ cells.

NGS library preparation was performed according to library’s user manual. shRNA library barcodes were amplified from genomic DNA using PCR and FwdHTS_R1 and RevHTS_R1 library primers. A second PCR reaction was performed from 5% of volume from the first PCR with nested library PCR primers (FwdGex_R2 and RevGex_R2) and 26-28 cycles. These primers include Illumina’s HiSeq adapter sequences. The nested PCR products were purified on a 2% agarose gel by gel-electrophoresis using the glass beads extraction method (48) and resuspended in 5 µl H2O. Next, the library was taken for NGS on the Illumina HiSeq2500 (Illumina, San Diego, CA, USA) using DECIPHER_GexSeqN sequence primer.

The shRNA sequencing results were initially analyzed using the DECIPHER specialized Barcode Analyzer and Deconvoluter software (Cellecta, Mountain View, CA, USA). The software yields a list of shRNA constructs and their sequencing read counts. First, the read counts were normalized to 20 million total sequencing reads. Next, for each shRNA construct the fold-enrichment between the norm.count in the GFP^high^ population compared to the control (GFP^low^ population) was calculated. A fold-enrichment threshold of 1.9 was set, and a gene was considered enriched if it had at least two shRNA constructs to pass the threshold. The fold enrichment of the gene was considered to be the maximal fold-enrichment of its shRNA constructs. Finally, a candidate list was comprised of genes that were found to be over 1.9-fold enriched in all three replicates. The genes were annotated using DAVID bioinformatics resources (49) for cellular localization and function.

### MafK constructs

For shRNA lentiviral vector generation, the best performing shRNA construct targeting MafK (shMafK) from the shRNA library (15) was used and shRNA double-strand oligos cloned into lentiviral pLKO.1-TRC-Puro cloning vector (Addgene plasmid #10878), according to The RNAi Consortium protocol..

shMafK_F ccggcacatggccaactaggatgaagttaatattcatagcttcatcctggttggccatgtgtttttg

shMafK_R aattcaaaaacacatggccaaccaggatgaagctatgaatattaacttcatcctagttggccatgtg

For MafK overexpression, MafK cDNA was PCR amplified and digested with BglII/EcoRI and cloned into pMSCV-Puro empty backbone (Clontech, Mountain View, CA, USA). For primers sequences used for cloning see **Error! Reference source not found.**.

### BAC IRF8 reporter constructs

The NIH3T3-BAC-IRF8.1 reporter cell-line was generated as previously described (14). To create BAC-IRF8.1-▽MafK-int6, the MafK-int6 binding-region on the BAC-IRF8.1 construct was replaced with a Zeocin antibiotic resistance gene under the regulation of the EM7 bacterial promoter, as previously described (14). The ▽MafK-int6 donor fragment was PCR amplified using primers containing 50 bps homology-arms adjacent to MafK-int6 binding-region, 5’arm_int6-em7 and 3’arm-Mafk_int6-zeo_R (see S1 Table). PCR reaction was done in LifePro Thermal cycler (Bioer, Hangzhou, China).) according to the following conditions: 94°C for 2 min, 40 cycles of 94°C for 30 sec, 50°C for 1 min and 64°C for 2 min, and finally 72°C for 7 min. PCR fragment was separated on 1% agarose gel-electrophoresis and purified using NucleoSpin Gel and PCR Clean-up (Macherey-Nagel).

### Generating BAC-IRF8.1-▽MafK-int6 stable clones

7.10^5^ NIH3T3 cells were seeded in each well of 6-wells tissue culture plates with 2 ml DMEM fresh media containing 5% FCS and incubated at 37°C in 5% CO2 incubator. 16-24 hrs later, the cells were transfected with 5 μg of BAC DNA using Metafectene-Pro (Biontext laboratories GmbH, Germany), according to manufacturer’s instructions. Cells were harvested 24 hrs later and plated on 10 cm culture dishes and after additional 16 hrs, Geneticin (G418) was added to select for stably transfected clones. Individual clones were isolated 14-18 days later, and the copy number of various transfected BAC-IRF8 reporter clones was determined by qPCR of isolated genomic DNA in comparison to endogenous single copy genes such as HPRT and Nramp1.

### Chromatin immunoprecipitation (ChIP)

Either 2.10^6^ or 10^7^ NIH3T3 cells (for histone PTM or MafK/Bach1 ChIP, respectively) were fixed in 1% Formaldehyde fixing buffer (50 mM HEPES pH 7.9, 100 mM NaCl, 1 mM EDTA, 0.5 mM EGTA, 1% Formaldehyde) and quenched with 125mM glycine. Chromatin was sheared using Covaris M220 Focus ultrasonicator (Covaris, Inc., Woburn, MA, USA) and so fixed cells were lysed according to Covaris manufacturer’s protocol. For the immunoprecipitation step, Protein-G-Dynabeads (Invitrogen, Carlsbad, CA, USA) were incubated with monoclonal antibodies, recognizing either specific histone PTMs, MafK or Bach1 for at least 8 hrs in 4°C. The following antibodies were used: αH3K27ac (Abcam, CAT#ab4729), αH3K27me3 (Upstate, CAT#17-622), αMafK (Santa Cruz, CAT#sc-22831 and anti-normal rabbit IgG (Upstate, CAT#17-622), αBach1 (C-20) (Santa Cruz, CAT#sc14700x). Sheared chromatin and coated-Dynabeads were incubated at 4°C over-night. Following chromatin-Dynabeads incubation, Dynabeads underwent several washes: twice with low-salt RIPA (10 mM Tris-HCl, pH 8.0, 1 mM EDTA, 150 mM NaCl, 1% Triton X-100, 0.1% SDS, 0.1% NaDOC), twice with high-salt RIPA buffer (10 mM Tris-HCl, pH 8.0, 1 mM EDTA, 500 mM NaCl, 1% Triton X-100, 0.1% SDS, 0.1% NaDOC),, twice with LiCl RIPA buffer (10 mM Tris-HCl, pH 8.0, 1 mM EDTA, 250 mM LiCl, 0.5% NP-40, 0.1% NaDOC) and once with TE. The Dynabeads underwent two elution steps, each with 100 µl ChIP elution buffer (10 mM Tris-HCl, pH 8.0, 5 mM EDTA, 300 mM NaCl, 1% SDS) and incubation at 65°C for 20 min. Eluates underwent reverse-crosslinking by incubation at 65°C over-night. Following reverse-crosslinking, samples were treated with RNase and Proteinase K (Sigma-Aldrich). ChIP DNA was purified using standard Phenol:Chloroform separation and Ethanol precipitation.

For histone PTM, ChIP samples were analyzed using real-time qPCR, as described hereafter (for primers sequences used see **Error! Reference source not found.**). For ChIP-Seq, samples were taken for NGS on the Illumina HiSeq 2500 (Illumina, San Diego, CA, USA). Library quality-control was performed using FASTQ software (version 0.11.5). For quality and adapter trimming trim_galore (cutadapt version 1.9) was used and sequencing reads were mapped using BWA aln v0.7.12. Peak calling was carried out by MACS2 v2.1.1 (20160309) and DiffBind v2.0.5 (R package) was used for peak merging and differential binding analysis (NIH3T3 binding compared to RAW binding). Finally, annotation were performed using ChIPseeker v1.8.3 (R package). For ChIP-PCR, ChIP samples were amplified using ReddyMix PCR Master Mix (Thermo Scientific, Waltham, MA, USA) with primers targeting MafK binding-region and resolved on 2% agarose gel.

### Real-time qPCR

The primers used for real-time qPCR were designed using PrimerExpress™ software (Applied Biosystems, Foster City, CA, USA), see **Error! Reference source not found.**. One µg of total RNA was reverse transcribed to cDNA using qScript™ cDNA Synthesis Kit (Quanta, Guishan District, Taiwan) according to manufacturer’s instructions. Reactions were performed using LifePro Thermal cycler (Bioer, Hangzhou, China). cDNA or genomic DNA was amplified using Power PerfeCTa SYBR^®^ Green SuperMix, Low ROX (Quanta, Guishan District, Taiwan) on the QuantStudio 12K Flex (Applied Biosystems by Life technology) according to the manufacturer’s instructions. Amplification conditions were as follows: 95°C for 20 sec followed by 40 cycles of 95°C for 1 sec and 60°C for 20 sec and then melt curve program of 95°C for 15 sec, 60°C for 1 min and 95°C for 15 sec. Relative gene expression level was determined using the dCt calculation method (gapdh gene was used as normalizer for cDNA amount and IRF8 2^nd^ intron was used as normalizer for genomic DNA).

### Luciferase reporter assay

Plasmid reporter constructs were generated by PCR amplifying MafK binding-regions (MafK25, MafK20 and MafK-int6) with primers flanked by MluI sites (see **Error! Reference source not found.**) and sub-cloned to pGL3 Luciferase vector (pGL3-Luc, Promega, Madison, WI, USA) driven by the IRF8 promoter (−1500 bps to the TSS). The MluI site is located upstream to the IRF8 promoter generating pGL3-Luc-MafK25, pGL3-Luc-MafK20 and pGL3-Luc-MafK-int6 reporter constructs. These plasmids were transfected to NIH3T3 or RAW cells and reporter-gene assays were performed 48 hrs later using the Dual-Luciferase Reporter Assay System (Promega, Madison, WI, USA), as previously described (14).

Retroviral reporter constructs were generated by PCR amplifying the pGL3 reporter cassettes described above and sub-cloning into the pMSCV retroviral vector, generating pMSCV-Luc, pMSCV-Luc-MafK25, pMSCV-Luc-MafK20 and pMSCV-Luc-MafK-int6, respectively (as illustrated in Fig. 5a). NIH3T3 or RAW cells were infected and reporter-gene assays were performed 72 hrs later (to ensure chromosomal integration) as previously described (14) using the Dual-Luciferase Reporter Assay System (Promega, Madison, WI, USA). For each sample, Luciferase light unit reads were normalized to genomic retroviral copy number, as determined using real-time QPCR with primers for Luciferase and IRF8 2^nd^ intron as reference (see **Error! Reference source not found.**).

### Flow cytometry

Flow cytometry analysis was performed using BD LSR-II flow cytometer (BD Bioscience, San Jose, CA, USA) and data was analyzed using Flowing Software 2 (Cell Imaging Core, Turku Centre for Biotechnology). For IRF8 staining, cells were fixed in 4% paraformaldehyde, permeabilized with 0.5% saponin, blocked with 10% normal donkey serum (Sigma-Aldrich) and stained with either goat anti-IRF8 (Santa Cruz, CAT# sc-6058) and CFL405 anti-goat IgG (Santa Cruz, CAT# sc-362245) or with CFL405 anti-goat IgG alone as control. Unstained wild-type NIH3T3 cells were used as negative control for GFP.

### Statistical methods

All Experiments were performed in n≥2 replicates and values are presented as means±AvDev. Data were compared by unpaired two-tailed Student’s t-test; p-values <0.05 or <0.01 were considered to be statistically significant, as indicated in the appropriate figure. When applicable False Discovery Rate (FDR) correction for multiple hypotheses testing was employed, using the Benjamini-Hochberg method (50). Asterisk indicates p-values that are significant after correction with α=0.05 or α=0.01.

## Data Availability

The MafK ChIP-seq datasets generated during the current study have been deposited in NCBI’s Gene Expression Omnibus (51) and is accessible through GEO Series accession number GSE113145 (https://www.ncbi.nlm.nih.gov/geo/query/acc.cgi?acc=GSE113145), respectively. The shRNA library screen datasets are available in S2 Table.

## Acknowledgements

We are grateful to Drs. Khateb, Koren and Barnea-Yizhar for critical reading of the manuscript, to the Technion’s Genome Center and the Russell Berrie Nanotechnology Institute for the support. This research was funded by The Israel Science Foundation [grant number 224/15 to B.Z.L]. B.Z.L. is an incumbent of the Lily and Silvian Marcus Chair in Life Sciences, Technion.

## Contributions

N.F. and B.Z.L. proposed and designed the experiments. N.F., M.Z., and A.A., performed the experiments. T.D. analyzed the barcoded shRNA library data. N.F. and B.Z.L. wrote the manuscript. B.Z.L. supervised the project.

## Supporting information

**S1 Fig. MafK binds IRF8 in human cell-lines.** Data tracks from ENCODE (22) depicting MafK ChIP-Seq data obtained from H1-hESC, HeLa, IMR90, K562 (IRF8-restrictive cells) and GM12878 (IRF8-permissive cells) covering human IRF8 locus (hg19). Data is presented using IGV genome browser (52, 53).

**S2 Fig. MafK ChIP-PCR validation of MafK binding in IRF8 locus.** Representative gels of MafK ChIP-PCR (Ig non-specific Ab is presented as negative control) with specific PCR primers targeting each MafK binding-regions (MafK25, MafK20 and MafK-int6), for validation of ChIP-seq results. Hmox1 primers are used as positive control.

**S3 Fig. MafK overexpression in NIH3T3 cells transfected with pGL3-IRF8p-Luc-MafK-int6.** NIH3T3 cells were transiently transfected with pGL3-IRF8p-Luc-MafK-int6 reporter construct and pMSCV-MafK. MafK expression level was measured using real-time qRT-PCR. Graph represents mean±AvDev of n=3.

**S4 Fig. MafK expression in different cell lines.** MafK relative expression was measured using real-time qRT-PCR in IRF8-restrictive cell lines (32D, NIH3T3) and IRF8-premissive cell line (RAW).

**S5 Fig. Bach1 involvement in the repressive activity of MafK regulatory regions in expression-restrictive NIH3T3 cells.** (a) NIH3T3 cells were transiently transfected with pGL3-IRF8p-Luc-MafK reporter constructs and pMSCV-empty (control) or pMSCV-Bach1 over-expression vector (Bach OE). Luciferase activity was normalized to Rennila activity and total protein amount. Relative Luciferase activity was calculated as ratio between MafK construct activity and Luciferase control construct (pGL3-IRF8-Luc empty vector). *p-value<0.05 student’s t-test, n=4. (b) Bach1 overexpression was validated using real-time qRT-PCR. Graph represents mean±AvDev of n=3.

**S6 Fig. Bach1 does not bind MafK regulatory-regions in IRF8 locus.** Representative gels of MafK and Bach1 ChIP-PCR (Ig non-specific Ab is presented as negative control) with specific PCR primers targeting each MafK binding-regions (MafK25, MafK20 and MafK-int6). Hmox1 primers are used as positive control.

**S1 Table. Primers list.** Sequences of all primers used in this study.

**S2 Table. ShRNA library screen.** Dataset of three replicates of the mouse DECIPHER pooled lentiviral shRNA library screen in IRF8-restrictive reporter cell-line NIH3T3-BACIRF8.1-GFP. Each library was normalized to 20M sequencing reads. For each shRNA construct fold change values were calculated as the ratio between GFP^high^ and GFP^low^ (control) read counts.

